# Fatigue and cognitive performance change in MS: multifactorial with disparate influences Short title: Predictors of fatigue and performance change in MS

**DOI:** 10.1101/414011

**Authors:** M Hu, N Muhlert, N Robertson, M Winter

## Abstract

**Background:** Fatigue is a common and disabling symptom in Multiple Sclerosis (MS) with a variety of direct and indirect influences, but remains poorly understood. Performance-based and self-report measures of fatigue are only weakly correlated and may have independent predictors. We adopted a multifactorial approach, utilising a measure of concurrent cognitive performance change in order to examine the clinical, psychological, and cognitive factors influencing subjective and objective fatigue in MS.

**Methods:** Sixty-one people with MS were assessed. Subjective fatigue was measured using the Modified Fatigue Impact Scale, Fatigue Assessment Instrument, and a Visual Analogue Scale (VAS). The Conners Continuous Performance Test 3 (CCPT3) and VAS were administered before and after two hours of cognitive testing, representing a period of cognitive effort. The differences in scores formed measures of objective performance fatigue and subjective fatigue change, respectively. We examined differences across baseline fatigue, fatigue change and performance change classifications, using regression analysis to uncover predictors of subjective fatigue and performance change.

Table 1.
Demographic and clinical features of the sample

Table 1. Demographic and clinical features of the sample
|  |  |
| --- | --- |
| **N** | 61 |
| **Age in Years** (mean [SD]) | 42.5 [11.3] |
| **Gender** (Female : Male) | 45: 16 |
| **Estimated general ability** (mean[SD]) | 109.3 [6.4] |
| **EDSS** (median) | 2.5 |
| **Duration from Disease Onset** (mean) | 10 years |
| **Duration from Diagnosis** (mean) | 5 years |
| **Relapsing Remitting** | 47 |
| **Secondary Progressive** | 9 |
| **Primary Progressive** | 5 |

**Results:** Depression, sleep, and emotion-focused coping each predicted baseline fatigue and together explained 53.5% of variance. Increased subjective fatigue was linked with anxiety, lower self-efficacy and gender. Cognitive performance change on the CCPT3 was however predicted by estimated general cognitive ability, self-efficacy and post-intervention fatigue.

**Conclusion:** Subjective fatigue in MS is a multifactorial construct, with subjective and objective cognitive performance fatigue largely influenced by indirect psychological and cognitive factors. The varying factors driving subjective and objective fatigue suggest that future studies need to take into account these disparate aspects when developing fatigue assessment tools. Targeting influential fatigue drivers such as psychological variables, and even using gender specific interventions may have the potential to improve the burden of fatigue and quality of life of people with MS.

## Introduction

Fatigue is common in multiple sclerosis (MS), with up to 92% of patients identifying it as one of the most prevalent and problematic symptoms^1^. It has been reported as gender invariant^2^ and has debilitating effects on physical function, activities of daily living (including employment and productivity), social relationships, psychological wellbeing, and quality of life^3^’^6^. The human and economic costs of fatigue indicate clear benefits in identifying and treating factors that contribute to its severity^4^. Unfortunately fatigue in MS remains poorly understood with management strategies that have been only partially effective at best^7^.

The variety of direct and indirect factors influencing MS fatigue has made it difficult to uncover predisposing factors^8^. Direct factors linking fatigue with biological disease characteristics include immune dysregulation and changes in brain structure and function^8,9^. Recent evidence also points to a causal role for proinflammatory cytokines, endocrine influences, axonal loss, and alterations in patterns of neural activity^10^. Fatigue is also indirectly related to factors such as sleep, pain, inactivity, mood, self-efficacy, medications, and other clinical variables^8,1113^. The understanding of MS related fatigue has been complicated by the interaction of these complex pathophysiological and psychosocial drivers.

There is a lack of consensus regarding a definition of fatigue^7,8,10^, which has led some studies to conceptualise fatigue as a unitary construct and others as a multifactorial symptom^8,14,15^. Fatigue in MS is typically considered to be the subjective lack of physical and/or mental energy that is perceived to interfere with activities^16^. An emphasis on subjective experience is reflected in the assessment of fatigue using self-report measures^17^. Unfortunately, these scales may not fully capture all aspects of fatigue, and tend to show weak associations with disease characteristics, objective performance change, and cognitive dysfunction^7,18^‘^21^. Subjective MS fatigue may be more closely associated with mood than neurological impairment^22^, therefore the need for concurrent measurement of objective fatigue is increasingly important.

Objective fatigue measures tend to focus on physical or physiological manifestations, but more recently have incorporated cognitive fatigue^7^. There is limited association between fatigue levels and physical disability as measured using Expanded Disability Status Scale (EDSS)^22,23^, highlighting an unclear relationship. Predictors of subjective and objective fatigue need to be analysed in greater detail, including their links with both physical and cognitive dysfunction^4,15^.

Our study asked whether subjective and objective fatigue in MS are differentially influenced by physical disability, psychological factors and cognitive function. We tested the hypothesis that subjective fatigue is predicted by indirect factors, such as mood, sleep and pain. In contrast, we hypothesised that objective measures of performance fatigue are predicted by physical disability and cognitive function.

## Methods

### Participants

A total of 61 participants were recruited from an established regional database of neuroinflammatory patients^24^. Inclusion criteria included; clinically definite diagnosis of MS^25^ within the last eight years; aged between 16-65 years old; and being fluent in English. The exclusion criteria included; history of other neurological or psychiatric condition; currently taking drugs known to substantially impact on cognition and/or fatigue (e.g. Baclofen); and having received a course of corticosteroids or disease modifying drugs within three months of recruitment. The study was approved by our local Ethics Committee (ref no. 05/WSE03/111).

### Measures and Design

The Modified Fatigue Impact Scale (MFIS), Fatigue Assessment Instrument (FAI) and Visual Analogue Scale (VAS) for fatigue provided three measures of baseline fatigue. The Conners Continuous Performance Test 3 (CCPT3), administered before and after intervention served as a measure of objective performance fatigue. The VAS was re-administered at the end to serve as a measure of subjective fatigue change. Each participant underwent an assessment of physical disability (EDSS) and completed a battery of psychological and cognitive measures (Figure 1). This reflected roughly 2.5 hours of cognitive effort.

**Figure 1.**
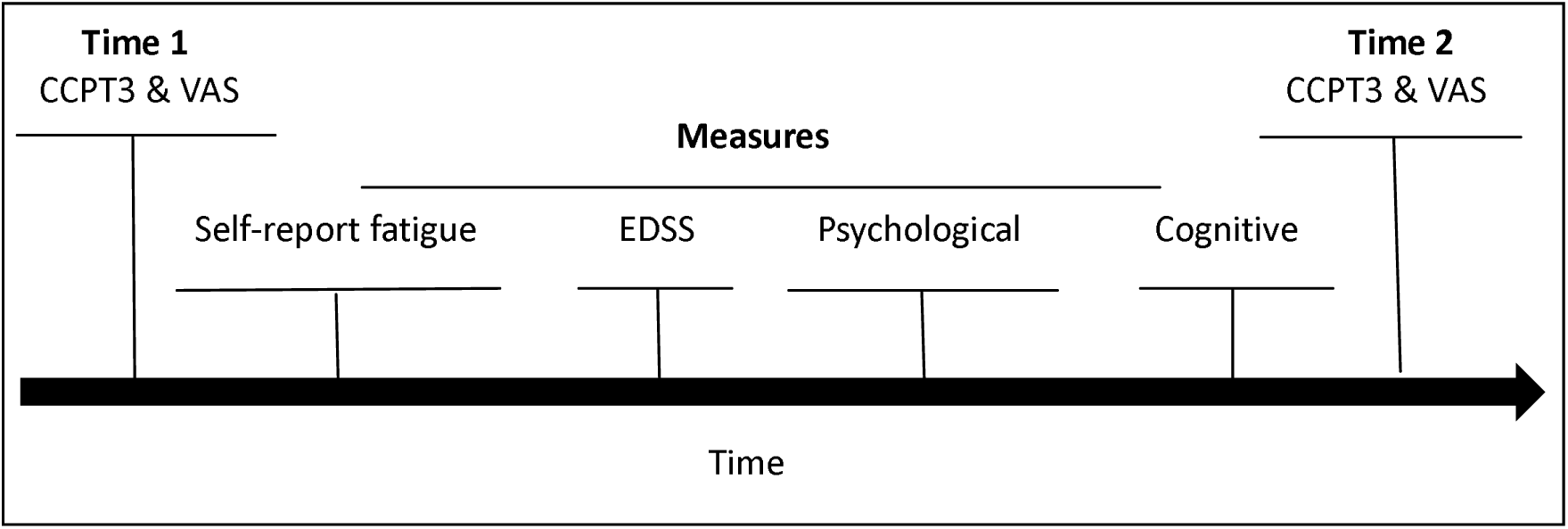
Administration procedure for all measures.

Psychological measures included scales of anxiety, depression, sleep, coping, pain, and self-efficacy. The cognitive battery comprised tests for estimating general cognitive ability, attention, learning, memory, information processing speed, motor speed, and executive functioning.

**Table 2.** Measures administered in between the CCPT3 sustained attention tasks. *The VAS was administered twice alongside the CCPT3*

| Physical | Fatigue | Psychological | Cognitive |
| --- | --- | --- | --- |
| EDSS | Modified Fatigue Impact Scale (MFIS) | Hospital Anxiety and Depression Scale | Wechsler Test of Adult Reading |
|  | Fatigue Assessment Instrument (FAI) | Medical Outcomes Survey Sleep Scale | Digit Span - Wechsler Memory Scale III |
|  | *Visual Analogue Fatigue Scale [VAS]* | Coping Inventory for Stressful Situations | BIRT Memory & Information Processing Battery |
|  |  | Pain Worksheet- Chronic Pain Coping Inventory | Delis-Kaplan Executive Function System<br>Letter & Category Fluency<br>Trail Making<br>Color-Word Interference<br>CWIT |
|  |  | General Self-Efficacy Scale | Alternate Uses Test |

### Statistical analyses

We used published cut-offs for the MFIS^25^ (i.e. 38) and the 11-item FAI Fatigue Severity Scale (FSS, i.e. 5)^26,27^to classify fatigue at baseline and ‘minimally important differences’ (MID)^28,29^ in pre-and post-intervention VAS scores to classify subjective fatigue change. Participants whose fatigue improved were grouped with those who remained stable, due to the small numbers. Performance change was determined by reliable change in pre-and post-intervention CCPT3 scores. The Reliable Change Index formula (CCPT3 manual) used standard error of difference to compute critical values. Group differences across classifications of fatigue, fatigue change, and performance change were examined for demographic, clinical, cognitive, psychological, and fatigue variables using independent samples t-tests and one-way between-groups analyses of covariance. Cognitive scores were converted to standard scores. We used the Chi Square Test for Independence with Yates Continuity Correction to examine differences across these classifications with gender as well as cognitive impairment status. We classified participants as cognitively impaired if two or more cognitive scores were at or below the 5^th^ percentile.

Using linear regression to predict baseline fatigue, the three fatigue scales used at the beginning of testing were reduced into a single fatigue factor using Principal Component Analysis (Figure 2).

**Figure 2.**
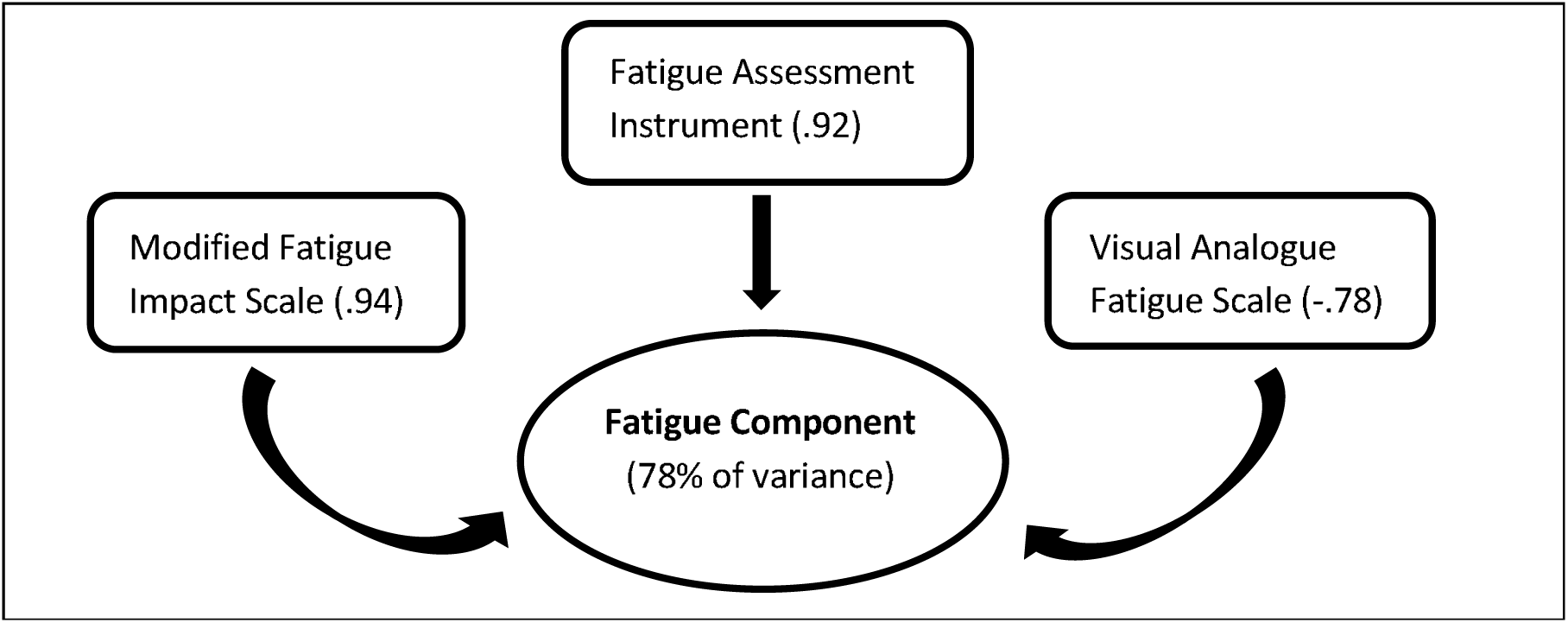
Principal component analysis showing the principal fatigue component (factor loadings shown in parentheses)

Inclusion of independent variables was informed by recommendations that these demonstrate bivariate correlations above .30 with the dependent variable, and less than .70 with each other^30^. Anxiety, depression, sleep, pain, coping (emotion focussed), self-efficacy, and EDSS were entered into the model with subjective fatigue as the dependent variable. Age, gender, disease duration, number of relapses, coping (task focussed and avoidance) and cognitive variables were excluded due to insufficient correlation.

Regression analysis was not used for fatigue change (difference between pre-and post-testing VAS scores) as this did not correlate with the fatigue factor or our other variables. Measures of performance change (differences between the first and second CCPT3 scores) showed insufficient correlations with most demographic, clinical, psychological and fatigue variables. The correlations meeting our criteria for linear regression were within the CCPT3 perseveration change and reaction time change variables (mean response speed and consistency of response speed). Estimated (general cognitive) ability and self-efficacy were entered into a model with perseveration change as the dependent variable. Letter fluency, CWIT Condition 3 (interference trial), number of impaired cognitive scores, and estimated ability were entered into a model with mean response speed change as the dependent variable. Post-intervention fatigue (2^nd^ VAS), estimated ability, visual learning, CWIT Condition 3, number of impaired cognitive scores, and avoidant coping were entered into the final model with response speed consistency change as the dependent variable.

## Results

### Baseline Fatigue

Roughly half of participants were fatigued at baseline using the MFIS or FAI with 39% classified as fatigued by both scales (Table 3). Those fatigued using the MFIS had greater fatigue on the baseline VAS (p=.005); more anxiety (p=<.0005) and depression (p=<.0005); poorer sleep quality (p=.015); greater pain (p=.007); more emotion-focussed coping (p=<.0005); less self-efficacy (p=.001); more disability (higher EDSS) (p=.008); and greater response variability (p=.011) on the first CCPT3 than those not fatigued. There was no association between MFIS classification and cognitive impairment status or gender.

**Table 3.** Fatigue classifications

|  | **MFIS**<br>**(cut off 38)** | **FAI FSS**<br>**(cut off 5)** | **Classification agreement** |
| --- | --- | --- | --- |
| **Fatigued** | **n=30 – 49%** | **n=28 – 46%** | **n=24 – 39%** |
| **Not fatigued** | **n=31 – 51%** | **n=33 – 54%** | **n=27 – 44%** |

The effect of MFIS classification on depression scores remained significant (p=.001) after the other variables demonstrating significant differences were controlled for. However, when adjusting for depression, only the differences in anxiety and emotion-focussed coping remained (p=.028 and p=<.0005, respectively).

Those fatigued using the FAI FSS had greater fatigue on the baseline VAS (p=<.0005); more anxiety (p=<.0005) and depression (p=<.0005); greater pain (p=.038); more emotion-focussed coping (p=.007); less self-efficacy (p=.025); higher EDSS (p=.018); and greater slowing of reaction times (p=.021) on the first CCPT3 than those not fatigued. There was no association between FAI FSS classification and cognitive impairment status or gender.

The effect of FAI FSS classification on depression scores remained significant (p=.014) when the other variables demonstrating significant differences were controlled for. However, after adjusting for depression, only the difference in emotion-focussed coping remained (p=.048).

For cognition, the group comparisons using the MFIS and FAI FSS cut-offs yielded a single difference (delayed visual recall, p=.023) with the former, and two differences (information processing, p=.03; motor speed, p=.044) with the latter scale. These effects disappeared with depression as a covariate. There were no differences in the number of impaired cognitive scores across classifications of either scale.

The linear regression model with the subjective fatigue factor as the dependent variable was significant (p<.0005) with 53.5% of the variance in subjective fatigue explained by the model (Adjusted R Square .535). Depression (p=.019), sleep quality (p=.017), and emotion-focussed coping (p=.014) made significant contributions to the variance in subjective fatigue. With shared variance partialled out, the unique proportions of variance accounted for by these variables were 4.5%, 4.6%, and 4.9%, respectively.

### Fatigue Change

Comparing pre-and post-intervention VAS scores, 35 (57.4%) rated their fatigue worse after intervention, 15 (24.6%) rated their fatigue the same, and 11 (18%) rated their fatigue as improved. Those fatigued at baseline (MFIS or FAI FSS) demonstrated greater post-intervention fatigue (p=.014 and p=<.0005, respectively) than those not fatigued.

However, the effect of MFIS classification on post-intervention fatigue disappeared when the variables demonstrating significant group differences at baseline were controlled for. The pre-intervention VAS alone accounted for a significant proportion of variance (37.6%) in post-intervention VAS fatigue (p=<.0005). Similarly, the effect of FAI FSS classification on post-intervention fatigue disappeared when the variables differing at baseline were controlled for. Unsurprisingly, pre-intervention VAS alone accounted for significant variance in post-intervention fatigue (VAS) (35.5%, p=<.0005).

Classification according to MIDs in fatigue resulted in fatigue worsening in 24 (39.3%) and either stable or improved fatigue in 37 (60.6%). Those whose fatigue worsened demonstrated more anxiety (p=.021), depression (p=.016) and less self-efficacy (p=.038), with no other differences across our variables. Depression means across groups were ‘normal’ (5.7 vs 3.3). The anxiety mean for those who worsened was ‘mild’ (8.7), and ‘normal’ for those who remained stable or improved (5.7).

There were no associations between the baseline fatigue classifications and fatigue change status (using MIDs). Similarly, there was no association between MID classification and gender, with 25% of males (n=4) and 44% of females (n=20) demonstrating worsened fatigue. However, grouping the raw fatigue change scores into ‘improved (or stable)’ and ‘worsened’ was associated (p=.03) with gender, with 31% of males (n=5) and 67% of females (n=30) demonstrating worsened fatigue. There were no gender differences in psychological variables, baseline fatigue measures, or post-intervention fatigue, but fatigue change (p=.044) differed. Females demonstrated more worsening than males, but this gender difference was attenuated (p=.055) once depression, anxiety and self-efficacy were accounted for.

Cognitively, those whose fatigue worsening was greater than the MID demonstrated more reliably changed CCPT3 scores (p=.002); greater worsening in reaction times (p=.038) and response speed consistency (p=.003); weaker visual learning (p=.039), information processing speed (p=.046), and category fluency (p=.004); as well as slower performance during the divided attention (Trail Making Condition 4, p=.015) and inhibition tasks (CWIT Condition 3, p=.008). After adjusting for estimated ability and the psychological variables that differed between groups, only the differences in number of reliably changed CCPT3 scores (p=.006) and response speed consistency change (p=.023) remained. There was no association between MID classification and cognitive impairment status.

When comparing the cognitively impaired (n=25, 41%) to those unimpaired, there were no differences in baseline fatigue variables, fatigue change or post-intervention fatigue. Whilst there were differences across cognitive variables, the number of reliably changed CCPT3 scores did not differ. There was a difference in EDSS scores (p=<.0005), with the impaired group demonstrating higher EDSS scores (mean [SD]= 4.3[2.2] versus 2.1[1.9]). The group effect remained significant (p=.002) accounting for 16.8% of the variance in EDSS scores with fatigue variables, depression, and age as covariates.

### Performance Change

We found 34 (55.7%) of our sample demonstrated reliable performance change on one or more CCPT3 variables (m=1.9, range 1-5) and 27 (44.3%) did not. Baseline fatigue variables did not differ between groups, but those with reliable change had more anxiety (p=.044), greater fatigue change (p=.006) and more post-intervention fatigue (p=.003). There was an association between fatigue change status based on MIDs and CCPT3 reliable change status (p=.001) with 38% of those whose fatigue remained stable or improved and 83% of those whose fatigue worsened demonstrating reliable change.

There were no differences on the baseline CCPT3, in estimated ability, or in the number of impaired cognitive scores. Whilst the reliable change group had slower motor speed (p=.03) and performance speed on the inhibition task (p=.013), there were no other cognitive differences. There were no differences in age, disease variables, EDSS, depression, sleep, pain, coping, or self-efficacy.

There was a significant association (p=.045) between reliable change status and gender, with 31% of males and 64% of females demonstrating reliable CCPT3 change. Females had significantly more reliably changed CCPT3 scores than males (p=.038), but they did not differ on initial CCPT3 scores. There was no association between gender and cognitive impairment status, and where the genders differed on cognitive variables (verbal learning, p=.001; information processing speed, p=.009; and motor speed, p=.021) females outperformed males. Males had longer disease duration (p=.048), but EDSS did not differ across genders.

### Predictors of performance change

The linear regression model with perseveration change as the dependent variable was significant (p=<.0005) explaining 24% of the variance in perseveration change (Adjusted R Square .24). Both estimated ability (p=.001) and self-efficacy (p=.005) made unique contributions to the model with little shared variance; 14% (Part Correlation .376) and 10.5% (Part Correlation -.325) respectively. The second model with reaction time change as the dependent variable was significant (p=<.0005) explaining 27.3% of the variance (Adjusted R Square .273). Estimated ability was the only independent variable to make a unique contribution (p=.037), which was only 5.5% of variance (Part Correlation .235) in reaction time change scores with shared variance partialled out. The last model with response speed consistency change as the dependent variable was also significant (p=.001) accounting for 24.5% of the variance in scores (Adjusted R Square .245). Post-intervention fatigue was the only independent variable to make a unique contribution (p=.014), accounting for 8% of variance (Part Correlation .283) in response speed consistency change scores with shared variance partialled out.

## Discussion

We identified that roughly half of our sample were classified as fatigued at baseline. Those who were fatigued were more likely to exhibit depression, anxiety, and emotion-focussed coping, and we found that depression, sleep quality, and emotion-focussed coping accounted for more than half of the variance in subjective baseline fatigue. These results appear to support our hypothesis of subjective fatigue being more strongly predicted by indirect factors than direct disease-related variables.

The links between mood and sleep and fatigue have been previously established^12^,^31^, and coping has been recognised as an important mediator between MS (including fatigue) and wellbeing^32^. Our results however suggest emotion-focussed coping has a direct influence on subjective fatigue. Whilst coping can predict depression in MS^33^, construct overlap cannot sufficiently explain our findings. We highlighted that whilst depression, sleep and coping may interrelate, they account for distinct contributions to subjective fatigue. Contrary to findings by others^34^ neither pain nor EDSS predicted our subjective fatigue component.

The pre-intervention VAS was the only baseline variable that accounted for a significant proportion of variance in post-intervention fatigue (2^nd^ VAS rating), but when those whose fatigue worsened (change greater than MIDs) were compared to those whose fatigue either improved or remained stable, there were no differences in baseline fatigue variables (including pre-intervention VAS fatigue ratings). There were no associations between baseline fatigue status and fatigue change status.

Those whose subjective fatigue worsened demonstrated more anxiety, depression, and less selfefficacy than those whose fatigue remained stable or improved. Whilst our gender results were mixed, fatigue change appeared to show little association with baseline fatigue, cognitive impairment, physical disability, or other demographic and clinical variables. The three psychological variables differing across groups suggest a role for indirect factors in both subjective baseline fatigue and fatigue change. Furthermore, this change was associated with reliable performance change and worsened performance consistency, in keeping with the possible effects of psychological variables on cognition^35-37^. Our association between fatigue change and performance change diverges somewhat from other studies reporting weak associations between subjective fatigue and objective cognitive performance change^38^. However, as baseline fatigue was not associated with fatigue change or performance change, and more than half the variance in baseline fatigue was accounted for by other variables, the role of fatigue as a driver for these changes is unclear.

We found no differences on baseline fatigue or pre-intervention CCPT3 variables between those who demonstrated reliable CCPT3 change and those who did not. However, those who demonstrated reliable change had more anxiety, fatigue change and post-intervention fatigue. There was an association between CCPT3 reliable change status and fatigue change status (based on MIDs) as well as gender. Whilst there was a link between cognitive impairment status and EDSS, neither had influence on subjective fatigue, fatigue change, or performance change. In our regressions, estimated ability made unique contributions to two performance change variables (perseveration and reaction time change). Self-efficacy also predicted perseveration change and post-intervention VAS predicted response speed consistency change. Whilst the results in performance change are mixed, our results provide little general support for the role of fatigue variables in performance change, particularly as most performance change variables (6/9) were excluded from further analyses due to insufficient correlations. Nevertheless, our second hypothesis gained limited support, but our results have shown that subjective fatigue and performance change may be influenced by different variables.

Fatigue in MS has often been found to be gender invariant^2^, but we found that gender featured in both fatigue change and performance change. Females demonstrated more worsening of fatigue and performance compared to males, despite not differing on psychological or fatigue variables (pre-or post-intervention), the first administration of the CCPT3, the number of impaired cognitive scores, or in cognitive impairment status. Where there were cognitive differences, females outperformed males. Once anxiety, depression and self-efficacy were adjusted for, the gender difference in fatigue change was attenuated. Anxiety and depression mean scores were all within the normal range and did not differ between genders (p >.2) with females demonstrating the higher mean for anxiety and males for depression. Males demonstrated higher mean (ie. better) self-efficacy (p=.07). Given the lower self-efficacy and higher anxiety scores in females, these two constructs may warrant further analysis alongside gender to determine their relative contributions to fatigue and performance change. If females with MS tend to experience more fatigue and performance change during prolonged cognitive effort, there may be implications for gender-specific intervention strategies. Interestingly, prolonged cognitive effort appeared to improve fatigue in 18% of our sample, suggesting a possible role for cognitive stimulation in improving subjective fatigue.

A limitation of this study is that we did not use a group of healthy controls. However, the validity of our results is supported by research into MIDs in fatigue^28,29^ and reliable performance change on the CCPT3. As part of the standardisation procedures this test was normed on 600 healthy adults (of which 384 covered the age range of our sample) with test-retest reliability measured on 63 adults with a mean age (43.5), similar to that of our sample. These norms may indeed enable more robust measurement of impairment and reliable change than using a small control group more vulnerable to sampling effects. Whilst using a cut-off to define cognitive impairment may aid study reproducibility, it cannot match the sensitivity or specificity of a bespoke clinical neuropsychological assessment in detecting impairment at the individual level^39^. It should be noted that our participants generally had low EDSS scores, and we did not differentiate between MS subtypes (majority relapsing remitting).

Future work could include designing fatigue scales with minimal construct and symptom overlap, with clear assessment of either subjective or objective fatigue. Measuring factors that may also contribute to the experience of fatigue and performance is important, so that relevant influences can be identified. Subjective fatigue (and even sustained performance to a degree) may be influenced by interventions for psychological variables such as depression, anxiety, coping, and self-efficacy. Providing targeted treatments have the potential to effectively enhance both psychological wellbeing and quality of life^30^.

We combined a computerised measure of objective cognitive performance fatigue with a multifactorial approach to fatigue assessment to highlight different factors in fatigue and performance change. In keeping with previous studies^8^, fatigue was not a unitary construct, and appeared more closely related to indirect than direct factors. Our results also suggest that predicting performance change is complex with aspects of cognitive performance influenced by different cognitive, psychological, and fatigue variables. In future we need to acknowledge multiple influences not only in examining subjective fatigue, but also when measuring cognitive performance change in relation to fatigue. Whilst there is still limited evidence-based fatigue management advice that can be offered^40^, there is an increasing drive to instigate multifactorial assessment and treatment of fatigue in MS^10^. Our study contributes to these efforts and we hope can promote discourse on the methods used to measure fatigue, and interventions best suited to reduce it.

## Acknowledgements

The authors wish to thank Dr Mark Wardle for his contribution to recruitment, which he enabled through a regional database. NM was funded by a Welsh Government Health & Care Research Wales fellowship (HF-14-21).

